# Long-run bacteria-phage coexistence dynamics under natural habitat conditions in an environmental biotechnology system

**DOI:** 10.1101/2020.04.21.053553

**Authors:** Leandro D. Guerrero, María V. Pérez, Esteban Orellana, Mariana Piuri, Cecilia Quiroga, Leonardo Erijman

## Abstract

Bacterial viruses are widespread and abundant across natural and engineered habitats. They influence ecosystem functioning through interactions with their hosts. Laboratory studies of phage-host pairs have advanced our understanding of phenotypic and genetic diversification in bacteria and phages. However, the dynamics of phage-host interactions has been seldom recorded in complex natural environments. We conducted an observational metagenomic study of the dynamics of interaction between *Gordonia* and their phages using a three-year data series of samples collected from a full-scale wastewater treatment plant. The aim was to obtain a comprehensive picture of the coevolution dynamics in naturally evolving populations at relatively high time resolution. Co-evolution was followed by monitoring changes over time in the CRISPR loci of *Gordonia* metagenome-assembled genome, and reciprocal changes in the viral genome. Genome-wide analysis indicated low strain variability of *Gordonia*, and almost clonal conservation of the trailer-end of the CRISPR loci. Incorporation of newer spacers gave rise to multiple coexisting bacterial populations. A host population containing a CRISPR array variant, which did not contain spacers against the coexisting phages, accounted for more than half of the total host abundance in the majority of samples. Phages genome co-evolved by introducing directional changes, with no preference for mutations within the protospacer and PAM regions. Metagenomic reconstruction of time-resolved variants of host and virus genomes revealed how selection operates at the population level. In activated sludge, it differed from the arms-race observed in nutrient rich media and resembled the fluctuating selection dynamics observed in natural environments.

## Introduction

Bacterial viruses (also bacteriophages or phages) are present in large number and are widely distributed in naturally functioning microbial ecosystems, including engineered environments, where they have a recognized key role in controlling microbial abundance and composition [1–3]. Well-designed experiments examining single host-phage pairs in the laboratory have advanced significantly our understanding of phenotypic and genetic diversification in bacteria and phages [4–6]. Longitudinal studies revealed that bacterial hosts under phage attack respond by evolving continuously, using a number of mechanisms to evade phage predation. In turn, genome rearrangements, mutations and antibacterial defense systems allow phages to overcome these barriers, leading to an evolutionary arms race [7–11].

However, the responses of bacterial cultures following phage infection in laboratory settings do not necessarily reflect the more complex interactions experienced in real world ecosystems, where bacterial hosts can be targeted by multiple phages [12], phages can infect more than one host [13], and density-dependent mechanisms may affect phage-bacteria coexistence [14] and immunity [15]. In addition, it has been recently demonstrated that bacteria-phage co-evolutionary interactions in natural environments are strongly affected by the complexity of biotic networks [16] and by the spatial structure of the environment [17].

Therefore, there is a strong need to complement lab-scale studies with field evaluation of the dynamics of phage-host interactions in naturally occurring microbial communities [18]. Even with the current advancement of metagenomics, monitoring the coevolutionary dynamics of phages and bacteria in the environment is not straightforward. Particularly challenging is the assignment of phages to their microbial hosts [19, 20]. Possibly one of the most suitable tool for the identification of phage–host pairs is the genomic region of clustered regularly interspaced short palindromic repeats (CRISPR), present in about half of bacterial and in most archaeal genomes. Within CRISPR arrays, bacteria and archaea accumulate short sequences of invading DNA, including phages, as a genetic memory of past invasions [21]. The presence of these foreign DNA sequences, named spacers, provides prokaryotes a molecular mechanism of immunity, which allows further defense against new attacks from the same entity [22]. Because a spacer sequence matches a protospacer sequence in the infecting phage, it can be used to pinpoint phage sequences within a complex metagenomic dataset and thus identify phage-host pairs. In addition, since new spacers are added at one end of the locus (the leader end), CRISPR loci provide a reverse chronological record of previous infections. The usefulness of this concept was demonstrated in a pioneer study, where phage and host microbial genome sequences were reconstructed from natural acidophilic biofilms [23]. Metagenomics was later successfully applied to address the role of CRISPR–Cas immunity in shaping microbial communities in a variety of natural environments [24–27].

There remain many unknowns about the dynamics of real populations of phages and bacterial in the environment [28]. A key challenge is to understand how phages and bacteria manage to coexist in complex natural environments, avoiding complete host and virus eradication [29]. To help bridge this gap, we conducted a detailed observational study of the dynamics of phage-bacteria interaction under natural habitat conditions, using a three-year data series of samples collected from a municipal full-scale activated sludge plant [30].

In this study, we focused on a bacterial host belonging to the genus *Gordonia*. Member of this genus have allegedly a prevailing role in biological foaming, a common operational problem affecting wastewater treatment plants (WWTPs) worldwide [31–34]. We used metagenomic data to reconstruct CRISPR-Cas loci of *Gordonia* populations. We then searched for corresponding protospacers in the metavirome, which allowed us to match phages to their specific bacterial host. The dynamics of phage-host interaction was followed by monitoring changes over time in the CRISPR loci of *Gordonia* populations, and the reciprocal changes in the viral genomes. The aim of the study was to obtain a comprehensive picture of phage-host coevolution in naturally evolving populations within an engineered environment at relatively high time resolution.

## Methods

### Wastewater treatment plant

Characteristics of the wastewater treatment plant, and full details of the sampling, nucleic acid extraction, and metagenomic analysis are provided in our previous article [30]. Briefly, samples were obtained from a full-scale municipal WWTP that provides sewage treatment for a population of ca. 600,000 residents of the metropolitan area of Buenos Aires (Argentina). A total of 60 samples of activated sludge collected biweekly over a period of 3 years from one aeration basin were stored at -80°C until analysis.

### DNA extraction and sequencing

DNA from sludge samples was extracted by a direct lysis procedure involving physical disruption of cells and a CTAB (cetyltrimethylammonium bromide) purification step [30]. The DNA extracted from the 60 sludge samples samples were sent to INDEAR (Rosario, Argentina) for Nextera DNA library preparation and sequencing. A rapid-run sequencing on two lanes was performed in a HiSeq 1500 Illumina, generating paired-end (PE) reads of 250 bp.

### *Gordonia* MAG assembly and characterization

Reads were recovered from the *Gordonia* MAG assembled in Perez 2019 [30]. We confirmed that all scaffolds belonged to the same MAG by calculating the abundance and tetranucleotide frequency correlation of each scaffold in the MAG. Relatedness of the MAG to known *Gordonia* species was evaluated on the basis of whole genome pairwise average nucleotide identity (ANI) [35].

The number and abundance of strains in the core *Gordonia* MAG that were present over the whole sampling period was inferred using DESMAN [36]. DESMAN pipeline was applied over marker genes extracted from checkM and filtered by COG annotation (only annotated genes), protein minimum length (330 aa) and total coverage (range 280X to 375X).

### Targeted metagenomic CRISPR array reconstruction

CRISPR-Cas system identification was done by detection and analysis of the cas operons using CRISPRCasFinder [37] and CRISPRDetect [38]. The leader regions were defined as the sequences spanning from the last *cas* gene to the first repeat.

After identified the repetitive sequence of each CRISPR, bbduck (bbmap software) [39] was used to extract all the reads from the metagenomic samples containing that sequence allowing 1 mismatch. These reads were used to reconstruct all the detectable variants of that particular CRISPR array as follow:

For each read from the previous step, the position of the repeat sequence was identified and used to trim the read into small sequences compose of seqL-repeat-seqR (s-r-s), where seqL and seqR were the left and right flanking sequences of the repeat. These s-r-s sequences were used as building blocks to reconstruct the CRISPR array network by pairwise alignment of the flanking sequences (s part of the sequence). If two s-r-s aligned by one of their extremes, both sequences were contiguous (s_1_-r-s_2_ + s_3_-r-s_4_ ==> s_1_-r-s_2=3_-r-s_4_). Therefore, if a sequence was flanked by repeats it had to be a spacer. The leader sequence was identified as the sequence that more often appeared as a “leaf” in the network. Its presence was later checked in the CRISPR-Cas locus of the *Gordonia* MAG. As a result, we obtained a network of all possible spacers (nodes) connected by repeats (edges). A series of parameters for nodes and edges, such as % mismatch, alignment length, coverage, sequence quality, were calculated and used to manually curate the network. For the analysis, a series of python scripts were created and used in combination with other available scripts from Mothur [40] (**Supplemental Fig S1**).

### Phage genomes assembly and annotation

The reconstructed CRISPR were used to predict phage-host associations [20]. We used the sequence of the CRISPR spacers to match phage protospacers in the metagenome obtained from the same longitudinal study. Matching contigs with identical temporal abundance profile were extracted and reassembled using Spades [41]. Base on this approach, two complete or nearly complete phage genomes were reconstructed. These genomes were used as templates to estimate their relative abundance mapping reads using bowtie2 [42] and the JGI script from Metabat2 [43]. When possible, reads from individual samples were assembled independently to identify potential genomics rearrangement.

Phages annotation was performed using DNAMaster [44] and curated manually.

Protospacers were identified in the phage genomes sequences using blastn, allowing for up to 5 mismatches, which were subsequently inspected manually. To identify the PAM sequence, 6 bp at both end of the protospacers were extracted, aligned and visualized using WebLogo [45].

### Analysis of sequence variants in phage DC-56 genome

Phage SNVs were identified in samples with a minimum coverage of 10X. The tool *Samtools mpileup* [46] was used to assess the proportion of each variant per position in the phage genome using the read mapping information (BAM files). Each nucleotide position was considered as an SNV when a mismatch to the consensus nucleotide at that position was supported for at less 5 reads. To reduce the possibility of including false positive SNV due to unspecific recruitment of reads, positions with coverage in excess of two-fold greater than the average coverage of the genome were excluded.

### Phylogenetic signal analysis

Differences in phage SNV composition among samples were identified using *bcftools gtcheck* [46] and represented as an error rates matrix. Phylogenetic signal analysis was performed with the R package phylosignal [47], using a cluster of samples based on the number of discordant SNV (error rate matrix) and time as trait. Random generated values and sample time randomization were used as control traits.

### Data availability

Sequencing data are available at NCBI BioProject under accession no. PRJNA484416. CRISPR reconstruction scripts are available at https://github.com/GuerreroCRISPR/Gordonia-CRISPR

## Results

### Virus assembly and annotation

Draft genomes of two phages, designated phage DC-56 and phage DS-92, were assembled each into one contig of 56 274 bp and 92 107 bp, respectively (**Supplemental Fig S2**). Structural variants between genomes assembled from individual samples were not detected. Therefore, reconstructed phage genomes were composite assemblies from all reads collected from samples in which phages were present.

G+C content of phages DC-56 and phage DS-92 were 65.2 and 52.4, respectively. Both phages belonged to the *Siphoviridae* family. Phage DC-56 was classified into cluster DC and phage DS-92 into cluster DS, according to the similarity with other *Gordonia* phages and members of the corresponding mycobacteriophage clusters [48, 49].

Phage DC-56 has 67 putative open reading frames (ORFs), 64 of which are transcribed in one direction and only three in the opposite direction (**Supplemental Fig S2A**). Like other *Gordonia* phages, phage DC-56 encodes two integrase genes, a tyrosine integrase (Int-Y, gene 46) and a serine integrase (Int-S, gene 47) near the center of the genome. Phage DS-92 encodes 111 potential ORFs, of which 46 are transcribed rightwards and 65 leftwards. We were not able to identify integrase genes in this genome.

### *Gordonia* metagenome-assembled genome

The *Gordonia* metagenome-assembled genome (MAG) was one of the 173 MAGs obtained in a previous study using data from a time-series analysis of a municipal full-scale activated sludge plant [30]. The refined *Gordonia* sp. MAG comprised 59 contigs, with a size of 5.1 Mbp, N50 of 158 040 and a G⍰+⍰C content of 66.35%. The analysis using single-copy marker genes [50] indicated that the assembly was 98.6% complete with less than 0.3% contamination. The MAG contained a total of 4 512 gene-coding regions, 58 tRNAs, and the complete sequences of 5S, 16S and 23S ribosomal RNAs. Therefore, the MAG complies with all standards of a high-quality draft [51].

Scores of genome pairwise average nucleotide identity computed with 87 genomes of recognized species of the genus *Gordonia* deposited in the NCBI database were <86%, well below the value of 95% that is considered the threshold for novel species definition (**Supplemental Table S1**).

Since *Gordonia* MAG was co-assembled using multiple samples across the time-series, it was important to assess the strain heterogeneity of the resulting assembly. Single-nucleotide variants (SNV) in 216 marker genes and co-occurrence across samples from the longitudinal study were used to investigate strain-level variations in the *Gordonia* MAG. Only 178 significant SNVs were distributed across *Gordonia* MAG, allowing the differentiation into four co-existing strains, with one strain dominating throughout the whole investigated period (**Supplemental Fig S3**). The very low genetic diversity between the four strains suggests that they derived from the same ancestor and most likely did not differ in their functional traits.

### CRISPR loci

*Gordonia* genome contained 2 CRISPR-Cas loci, classified as type I-E and type III-B-like systems, based on the architecture of the Cas operon (**Supplemental Fig S4**). Automatic prediction of CRISPR arrays from MAGs may underestimate the diversity of spacers because of the limited capacity to resolve the multiple CRISPR variants present in a mixed population. To overcome this problem we used a specific pipeline to reconstruct all observed CRISPR variants, based on the overlapping of every possible connection between spacers identified in raw reads across the complete time-resolved metagenomic dataset (see M&M) [52–54]. The reconstruction of the type I-E CRISPR array, designated CRISPR-1, can be visualized in the network shown in **Fig 1**. The spacer arrangement of CRISPR-1 showed one particular spacer (node circled with a thick line in **Fig 1**) acting as a breaking point, after which the diversity of CRISPR variants increased markedly in the direction of the leader end. Thus, spacers were highly conserved in the direction of the trailer-end of the CRISPR locus, whereas new spacer acquisition led to a considerable increase in the CRISPR population.

**Fig 1:**
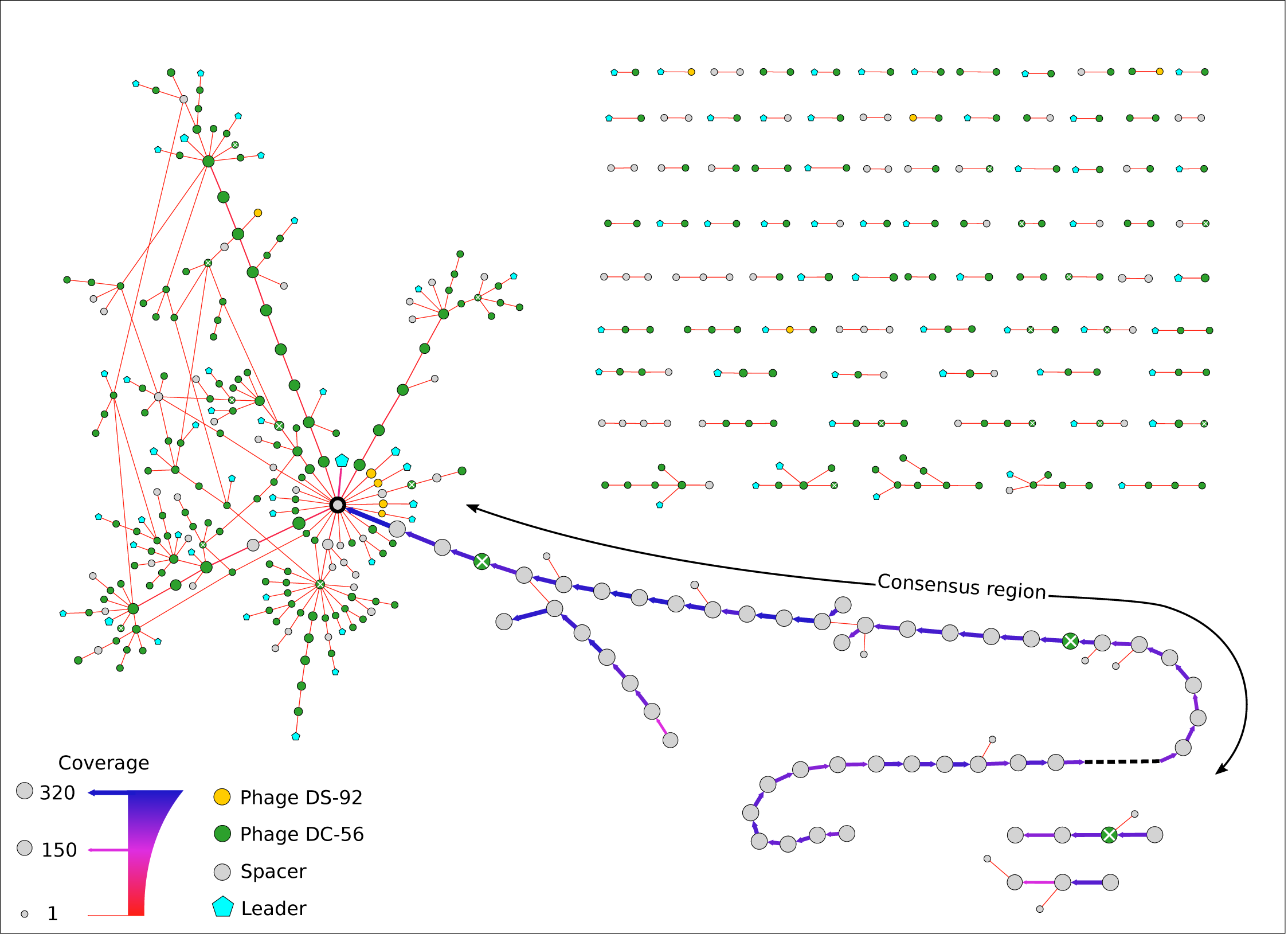
Reconstruction of the spacer arrangement of *Gordonia’s* CRISPR-1 across the complete set of metagenomes from Perez et al. [30]. Each circle represents a spacer. Leader sequence are shown as pentagons. The lines connecting each spacer represent their position relative to other spacers, with arrows pointing to the direction of the leader sequence. Coverage is indicated by the diameter of the nodes and the width and color of the lines. Every combination of nodes connected by edges pointing in the same direction represents a possible CRISPR variant in the population. Spacers targeting phage DS-92 and phage DC-56 are colored with yellow and green, respectively. Spacer with mismatches with phages’ protospacers are marked with an X.

The low-coverage, fan-like-structure at the leader end (>=1X, **Fig 1**) confirmed the active incorporation of new spacers. The majority of these newly acquired spacers perfectly matched sequences of one of the coexisting phages (DC-56). Few spacers also matched sequences of the phage DS-92 genome, notably in the short-branched variants, suggesting simultaneous adaptation to multiple phages. The proximal end of most CRISPR-1 variants contained leader sequences, which are essential for the acquisition of new spacers. Spacers within the consensus region did not match phage DC-56 genome, and no leader sequences could be detected downstream of the breaking point node.

The average coverage of spacers within the consensus was similar to the average coverage of contigs from the complete *Gordonia* MAG, suggesting that the majority of *Gordonia* cells carried the CRISPR-1 array (**Supplemental Fig S5**). Upon filtering the low coverage nodes, 11 main variants of the CRISPR-1 were identified upstream the breaking point node (**Fig 2A**). Most of the spacers within these main branches matched to the co-existing phage DC-56, but four of them showed 1 and 2 mismatches (**Fig 1**). The abundance distribution of the main variants over time is shown in Fig 2B. The CRISPR-1 variant with a leader sequence immediately upstream the consensus region (named L1 in **Fig 2**) accounted for more than half of the total abundance in the majority of samples (**Fig 2C**).

**Fig 2:**
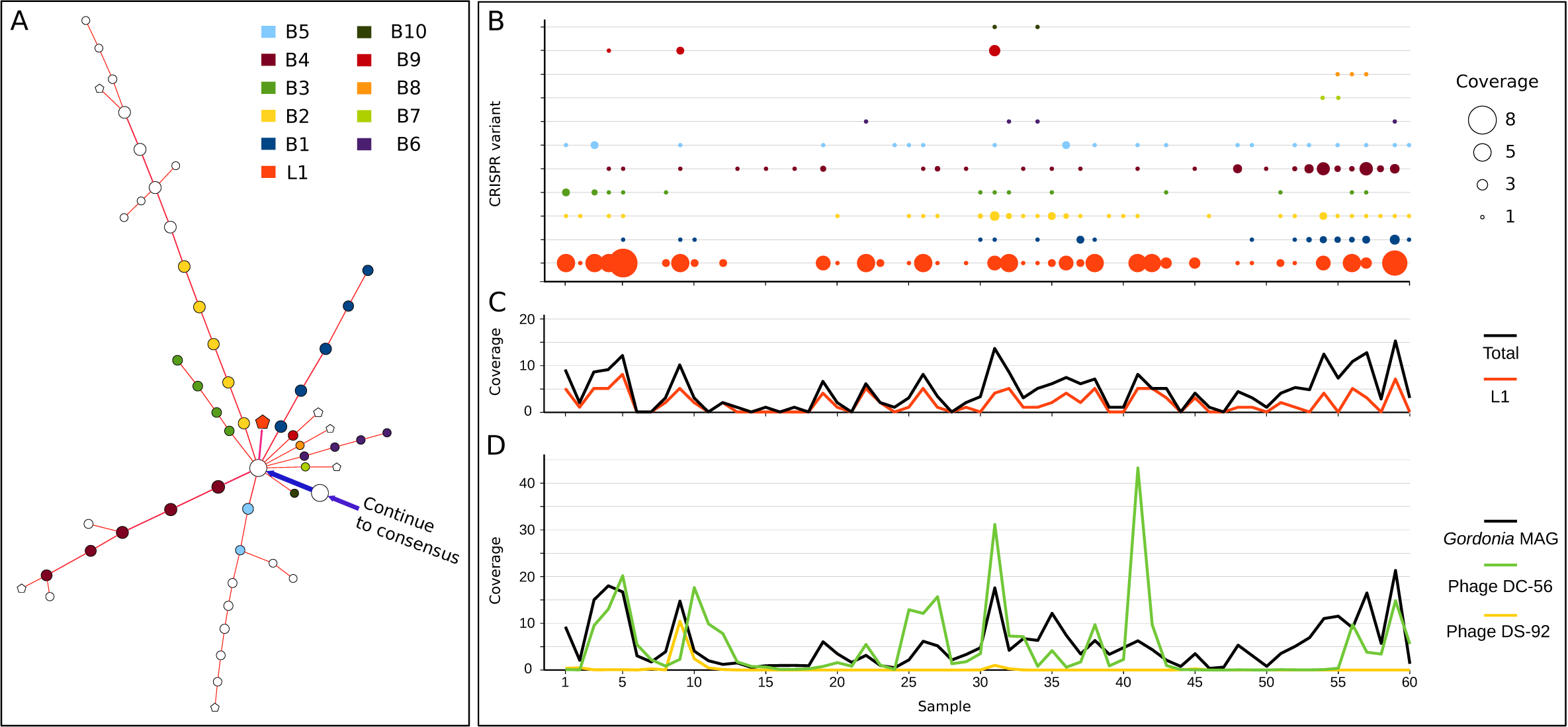
Abundance of the main CRISPR-1 variants over time. A) Simplified representation of Fig 1 showing nodes with coverage > 1 of the main variants. Nodes used to estimate the average abundance of each variant are identified by colors. B) Abundance of each variant per sample. C) Sum of all variants in panel B compared to variant L1. D) Time course of the abundance of the *Gordonia* MAG and the phages DC-56 and DS-92.

The reconstructed type III CRISPR locus, designated CRISPR-2, contained only 8 spacers (**Supplemental fig S4, Supplemental fig S6A**). The fact that the coverage of this locus matched the coverage of the *Gordonia* MAG (**Supplemental fig S6B**) indicates that it was likely also present in most members of the *Gordonia* population. Nevertheless, the lack of spacers matching coexisting phages and the absence of “branching” suggested that this system was either inactive or at least was not actively incorporating new spacers during the time span of this study.

### Protospacer adjacent motif sequences

The protospacer adjacent motif (PAM) sequence was searched by alignment of short stretches of sequences (up to 6 bp) immediately flanking the protospacer regions of phage DC-56 sequence. The trinucleotide GTT was highly conserved immediately at the 5Lr end of those protospacers that matched spacers in CRISPR-1 (**Supplemental fig S7**), which suggests that this is the PAM sequence required for incorporation of new spacers [55].

### Phage-host dynamics

Time course of phages and *Gordonia* coexistence is shown in **Fig 2D**. There was considerably overlap in peaks of abundance of bacteria and phage DC-56 over the time of the study (Pearson correlation r=0.42, p=0.0008; **Supplemental Table S2**), suggesting that bacteria have evolved towards phage resistance. At the same time, it is inferred that phage replication and survival was allowed by the presence of populations of susceptible bacterial cells. This is supported by the fact that peaks of phage DC-56 coincided especially with time points of higher abundance of *Gordonia* bacterial cells lacking spacers against the virus (**L1, Fig 2** Pearson correlation r=0.50; p=4.10^−5^; **Supplemental Table S2**). Phage DS-92 peaked with different abundance at two different times, in which *Gordonia* was also present in high abundance (**Fig 2D**).

### Adaptation of phage DC-56

To analyze the viral response to CRISPR-1 diversification, we focused on the sequence of phage DC-56 genome, whose abundance mirrored *Gordonia’s* MAG. The 285 spacers that targeted the phage DC-56 genome were distributed across the entire phage genome (**Fig 3**). There is an indication of a moderate spacer acquisition bias, although we found no evidence that the location and strand specificity of protospacers were influenced by previously acquired spacers. We also could not relate genomic regions with less density of protospacers with the presence of higher frequency of the canonical *E. coli* Chi sequence [56]. Nevertheless, with the available data we cannot ruled out that low protospacer density regions are protected by a different Chi sequence that is recognized by Gordonia’s RecBCD system, or the involvement of other processes such as interference-driven or primed spacer acquisition [57].

**Fig 3:**
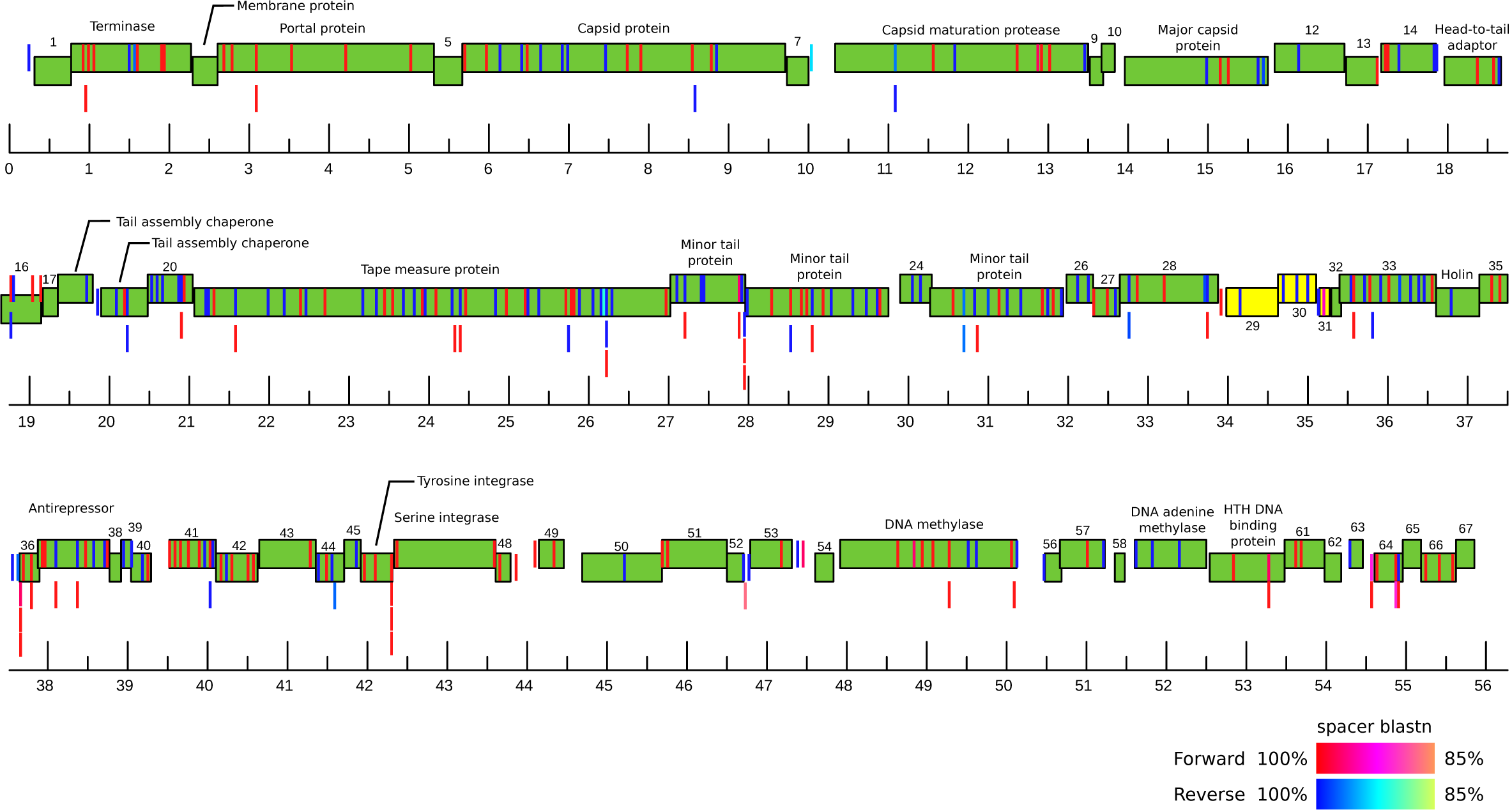
Phage DC-56 genome map highlighting the position of protospacers. Genes are indicated by boxes, colored according to their forward (green) or reverse (yellow) orientation. When known, the name of the genes was added. The location of protospacers targeted by CRISPR-1 was indicated by verticals bars. Bar colors indicate the orientation and the similarity to the corresponding spacer sequence in the *Gordonia* MAG.

Changes in phage DC-56 genome over three years were investigated through viral SNV analysis. Each nucleotide position was marked as an SNV when a mismatch to the consensus nucleotide at that position had a minimum coverage of 5. A total of 610 SNVs satisfied this criterion. To test for differences in the efficacy of variants selection, we examined the ratios of nonsynonymous to synonymous polymorphisms in protospacers and elsewhere in the phage genome and found that the ratio was not significantly different (two-tailed Fisher’s exact test P =0.66). Eighty-two SNVs were located within 54 protospacers and 5 PAM regions (**Fig 4**). Fisher’s exact test revealed that SNVs do not occur in protospacers more frequently than in the rest of the genome (p=0.11).

**Fig 4:**
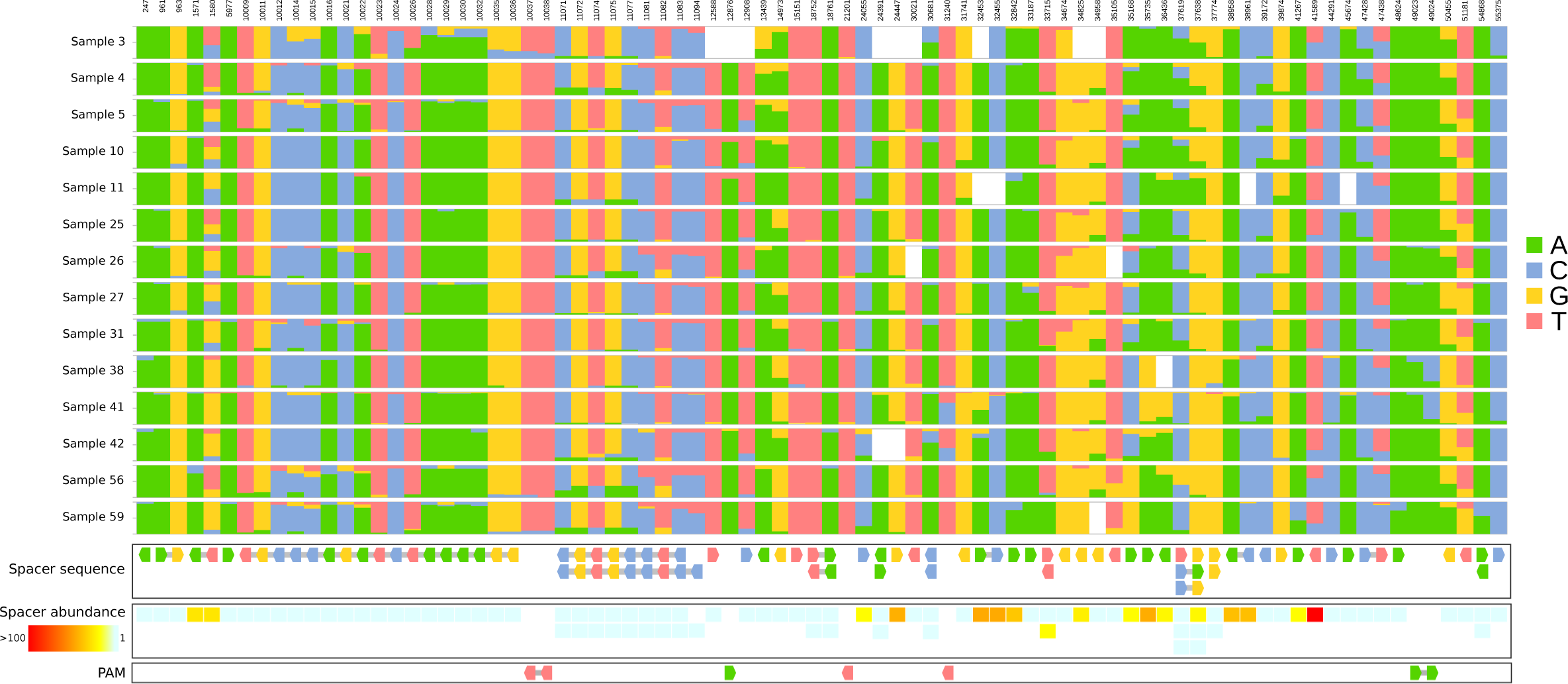
The 82 single-nucleotide variants (SNV) of phage DC-56 affecting protospacers and PAM regions. Only samples with a total coverage >=10 and a minimum coverage of 5 reads per sample were analyzed. Spacers are drawn as arrows to indicate orientation. Bases within the same spacer are connected by a gray bar. Bases in spacers located in the reverse orientation were colored according to the corresponding forward sequence.

Phylogenetic signal was used to investigate the pattern of longitudinal population dynamics of phage DC-56. We looked at the relationship between DC-56 sequence in each sample and time [58]. The significant temporal autocorrelation between the DC-56 phage population point to the higher similarity of samples closer in time than samples separated by longer time intervals, an indication of a directional trend in the variation of phage DC-56 genome sequence (**Fig 5, Supplemental Fig S8**).

**Fig 5:**
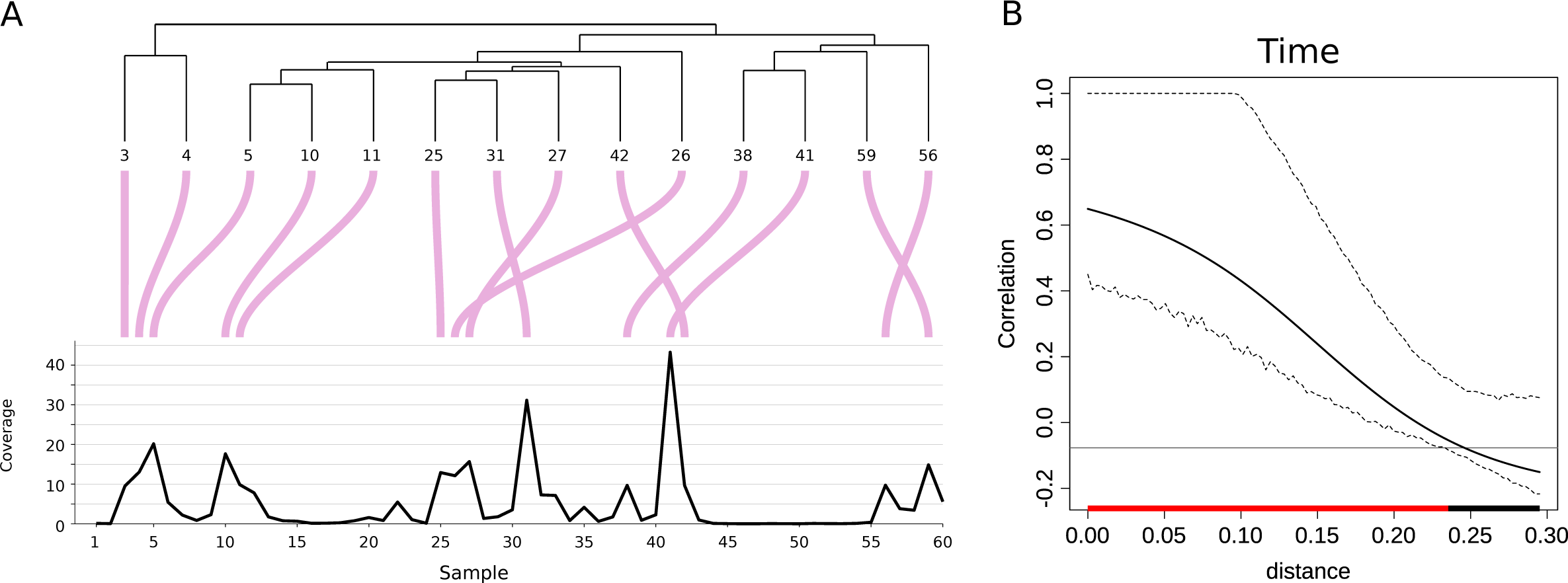
Phylogenetic signal analysis of reconstructed phage DC-56. A) Cluster of phage DC-56 genome based on shared SNV in samples with coverage >= 10 and their correspondence to phage abundance profile (coverage) over time. B) Correlogram of the distance between samples using time as trait. Red bar indicates positive correlation.

## Discussion

This work provides a comprehensive picture of the complex dynamics of naturally evolving populations of a bacterial host and its infecting phage within an engineered environment. Metagenomic reconstruction of time-resolved variants of host and virus genomes revealed how selection operates at the population level. Diversification of the bacterial CRISPR locus, which occurs as a result of phage infection, gives rise to multiple coexisting populations of a *Gordonia* strain. The majority of newly acquired spacers matched sequences of one of the coexisting phages. These observations coincide with simulations [59], experiments performed under laboratory settings [60, 61] and observation from natural environments [27], confirming that CRISPR array evolution is driven by co-occurring phages and suggesting that the synergistic effect of spacer diversity established at the host population level aim at reducing the probability of phages to escape mutations [60, 61].

### Phage-host dynamics

Under the arms race dynamics, phages would co-evolve to escape CRISPR by preferentially selecting variants with polymorphic positions at protospacer regions. For example, directional changes in the phage genome were observed upon long-term coincubation of *S. thermophilus* DGCC7710 and phage 2972 in milk medium [10]. This effect, which has also been observed in other phages, helps them to overcome sequence-specific CRISPR immunity [62, 63]. In contrast, in our study we observed directional changes, yet accumulation of mutations in the PAM and protospacer regions was not favored (**Fig 4**). This is not surprising, given that phages that escape bacterial immunity by mutating their target protospacers within one host will not be successful against coexisting bacterial populations that possess different spacers [64]. Yet, as long as no individual bacterial population achieves a much higher density than their competitors, phages would still find excess of susceptible hosts, reducing the selection pressure to mutate their targeted protospacers or PAM regions. As a matter of fact, the most abundant *Gordonia* population observed throughout this study contained a CRISPR array with no spacers matching protospacer sequences of phage DC-56 (L1 in **Fig 2**). The continuous presence of a large population of susceptible host may act as a propagating host for the phage DC-56, granting its endurance [11, 65]. Nevertheless, we cannot tell whether this is a non-immune population, associated with the fitness cost of maintaining a fully active CRISPR-Cas system [66, 67] or that the population maintains active different defense mechanisms or it is protected from phage exposure by the spatial heterogeneity of the environment [68].

It follows that in the long run neither the host nor the virus are driven extinct, in opposition to the results of laboratory coevolution experiments, where spacer diversity of CRISPR-mediated resistance caused rapid phage extinction [60]. Rather than a typical unidirectional arms race dynamics, the outcome of phage-*Gordonia* coevolution in activated sludge most closely resembled a fluctuating selection dynamics (FSD) [69]. FSD has been reported to occur in environments with low resource availability, such as soil [70] and the ocean [71], where the costs of phage resistance are expected to be high [72–74]. This is not entirely unexpected considering that *Gordonia* thrives in activated sludge, where bacteria grow under carbon limitation conditions, with typical loading rates in the range of 0.05 to 0.4 kg Biochemical Oxygen Demand per kg dry biomass and per day [30, 75, 76]. Yet, we cannot confirm whether the fluctuation of phage and bacterial genotypes is associated to a negative frequency-dependent selection of virus and hosts with different infectivity and resistance profiles [18].

CRISPR-Cas systems are one among several mechanisms that have evolved in bacteria to fight phages [73, 77]. In its natural habitat, *Gordonia* would be protected by several lines of defense, including the protecting effect of EPS and spatial heterogeneity of the matrix. However, the fact that phage DC-56 appears to evolve relatively slowly in this environment suggests that it is the continuous presence of large number of susceptible cells that makes it possible to maintain the stable phage-host coexistence.

A potential limitation of this study is that we cannot rule out the existence of additional hosts of the phages. Nevertheless, the dynamics of *Gordonia*-DC-56 phage coexistence shows a pattern in which the most abundant phages replicate actively as long as the host populations remain susceptible to their attack, as predicted by the “killing the winner” model [78]. The appearance of peaks of infection by a second phage are in turn consistent with the seed bank model, which postulates that minor virus populations are maintained in a bank fractions and can potentially become active when the environment changes [13].

### Conservation of trailer-end spacers

This work confirms earlier observations that spacers at the trailer-end of the CRISPR locus is highly conserved despite the fact that they would not provide immunity against coexisting viruses [24, 79, 80]. Simulations have predicted that trailer-ends may become clonal across time due to selective sweeps of highly immune CRISPR lineages [81]. More generally, the purge of genetic heterogeneity at the trailer-end of the CRISPR locus appears to conform to the genome-wide sweep predicted by the ecotype model [82], where a single population (ecotype) acquires an advantageous trait that helps it to survive, reducing overall population diversity. The occurrence of genome-wide selective sweeps has already been revealed in freshwater lake microbial communities, where with few exceptions, overall genetic heterogeneity of most bacterial populations was relatively low over a period of several years [83]. Nonetheless, the low bacterial strain diversity and the existence of a conserved trailer-end is puzzling because it suggests that the entire *Gordonia* population observed over 3 years appears to share a common ancestry, despite the fact that *Gordonia* are widespread in the environment and activated sludge is an open ecosystem, in which immigration plays an important role in community assembly [84–86].

### Implications for the application of phage therapy in environmental processes

*Gordonia* belongs to a group of Actinobacteria that have relatively high content of mycolic acids on their cell surface, called thereafter Mycolata. The highly hydrophobic cell surface of Mycolata in combination with their ability to excrete extracellular polymers, which may act as biosurfactants, favor the formation and stabilization of foams in wastewater treatment processes [87, 88]. Despite major efforts to find universal strategies to prevent or to control the unwanted proliferation of these bacteria, foaming remains a common operational problem in activated sludge. In recent years, considerable research has been directed towards the use of bacteriophages as an alternative approach to chemicals to reduce the levels of filamentous bacteria in wastewater treatment, including foam forming *Gordonia* [34, 88, 89]. The results obtained in this work suggest that the presence of an active lytic phage might not be sufficient to control *Gordonia* abundance. Successful application of phage therapy at the full-scale will still require a better understanding of virus-host interactions in the complex real-world context.

### Conclusions

In activated sludge, the pattern of coevolution of *Gordonia* and its phages differed from the arms-race dynamic observed in nutrient rich media and resembled the fluctuating selection dynamics observed in other natural environments with low resource availability. This hypothesis can be tested in further investigation on phage infectivity and host range and bacterial resistance [5, 27], as well as the protection effect due to spatial refuges for susceptible bacteria [90].

More than fifteen years ago, there was a call proposing wastewater treatment as model systems to address fundamental ecological questions in realistic environmental conditions [91, 92]. We renew this call, proposing that wastewater treatment, in conjunction with metagenomic time-series studies, are suitable means to gain further insight into bacteria-phage dynamics and interaction in real populations.

## Acknowledgments

This work was supported by grants from AySA-CONICET (Res. 3816/11 and 1371/15) and FONCyT (PICT 0746/15). The funders had no role in study design, data analysis, decision to publish, or preparation of the manuscript.

The authors declare no competing financial interests.

## Legends to Supplemental tables and Figs

**Supplemental table S1.** Pairwise ANI of *Gordonia* metagenome assembled genome compared to all available *Gordonia* genomes on NCBI (at March 2020)

**Supplemental Table S2**. Pearson correlation between the abundance of phage DC-56 and the abundance of the different CRISPR-1 variants over the time span of the study.

**Fig S1**. Schematic representation of the pipeline used to reconstruct the CRISPR arrays. A detailed explanation and scripts used can be found at https://github.com/GuerreroCRISPR/Gordonia-CRISPR.

**Fig S2**. Genome maps of *Gordonia* phages DC-56 (A) and DS-92 (B). Genes are indicated by boxes colored by orientation (green and red, for forward and reverse genes respectively). Ruler scale: 1000 bp.

**Fig S3**. Abundance of co-existing *Gordonia* strains over time.

**Fig S4.** CRISPR-Cas systems from *Gordonia* sp. A) Type I-E system. B) Type III-B-like system. Repeats are depicted as gray diamonds (Rn) and spacers as small vertical rectangles (Sn). Horizontal arrows represent the Cas operon, empty arrows represent putative RAMP genes and small subunit cas11. RT represents the group II introns reverse transcriptase.

**Fig S5**. Average coverage of the spacers within the consensus CRISPR-1 arrangement in *Gordonia* MAG (red line) and total *Gordonia* MAG (blue line) over time.

**Fig S6.** A) Reconstruction of the spacer arrangement of Gordonia’s CRISPR-2. B) Average coverage of the spacers within the CRISPR-2 arrangement in *Gordonia* MAG (orange line) and total *Gordonia* MAG (blue line) over time.

**Fig S7.** WebLogo for the PAM consensus sequence predicted for phage DC-56 required for CRISPR-1.

**Fig S8**. Phylogenetic signal analysis for phage DC-56 genome over time. Randomized time and random generated values were used as control traits. A) Data traits visualization mapped along the samples cluster. B) Correlograms for traits. Red bar indicates positive correlation.

