## Supplementary figures and images for "Long-run bacteria-phage coexistence dynamics under natural habitat conditions in an environmental biotechnology system"

### Supplementary Fig S1

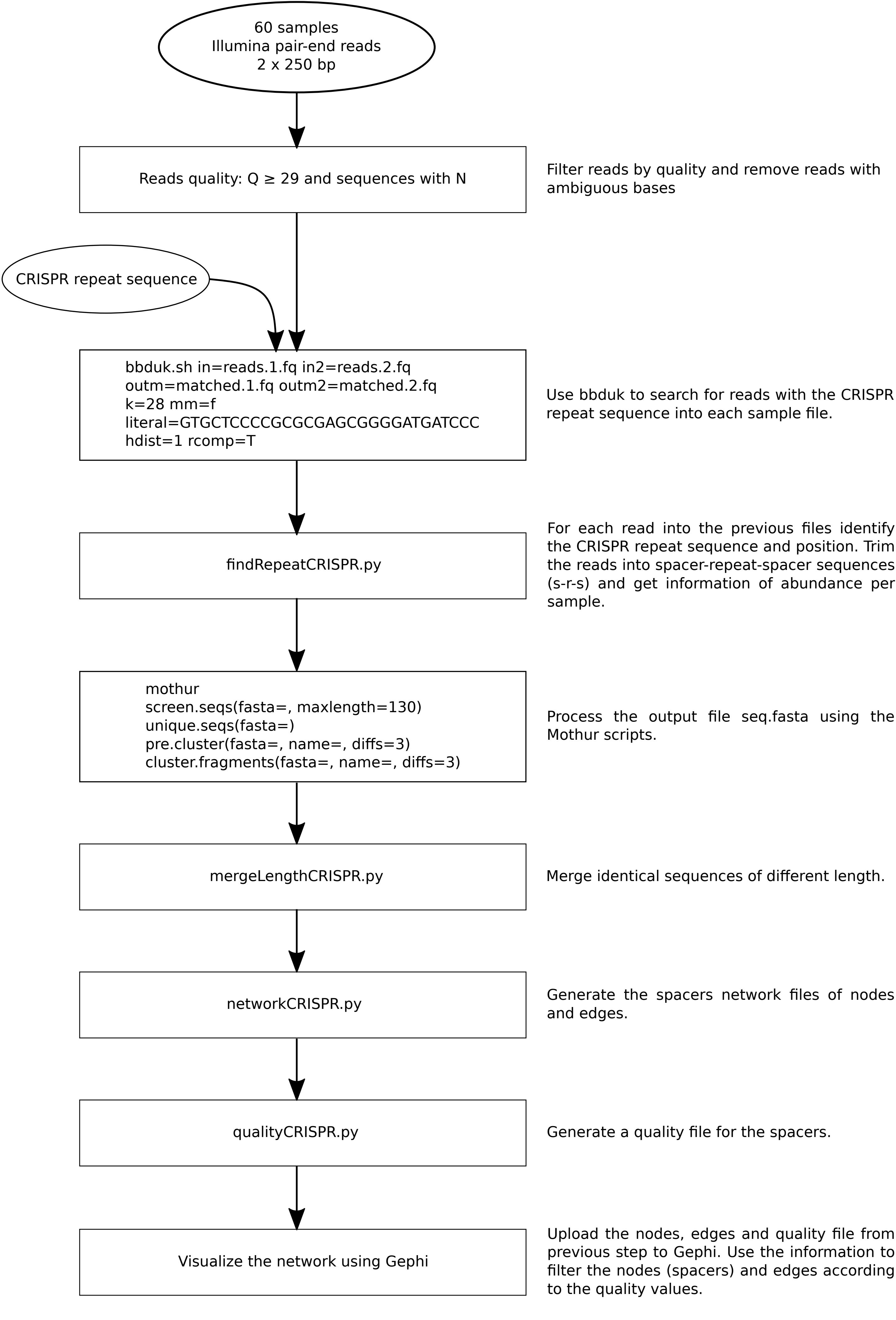


Supplementary figure S1

### Supplementary Fig S2

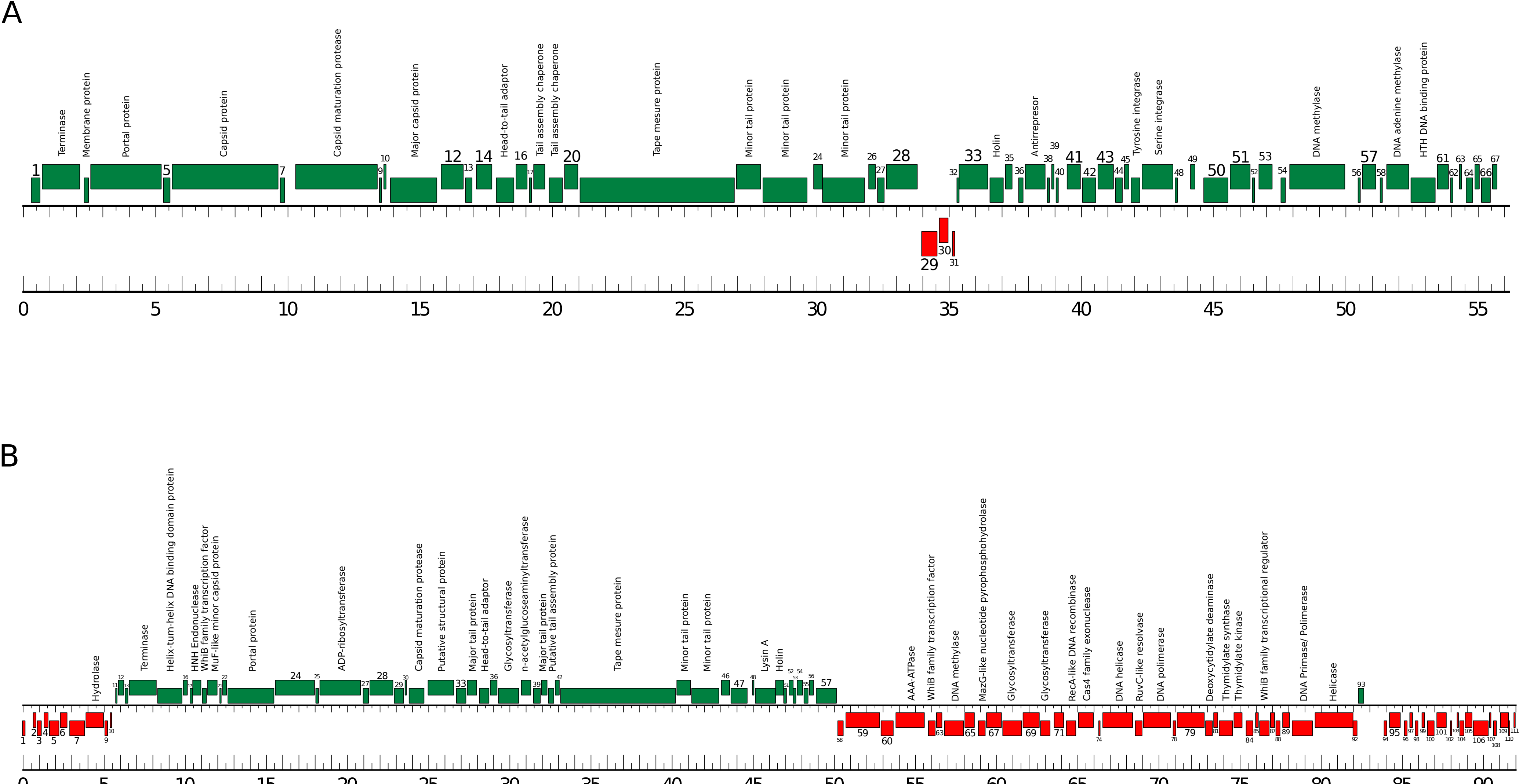


Supplementary figure S2

### Supplementary Fig S3

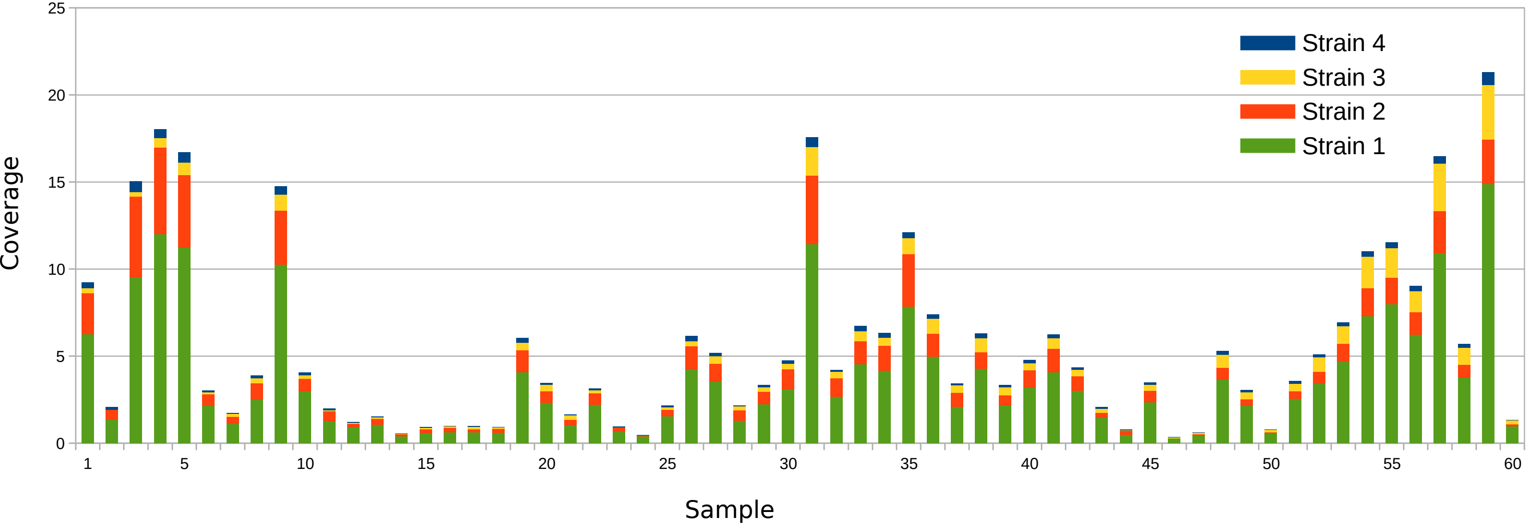


Supplementary figure S3

### Supplementary Fig S4

A

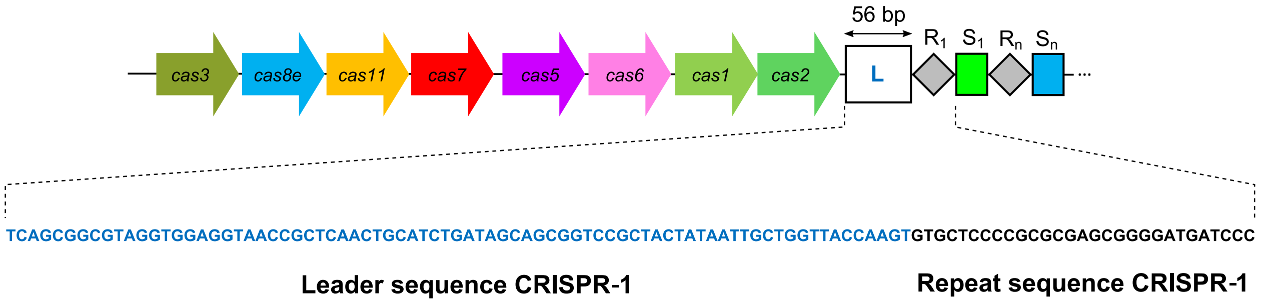


B

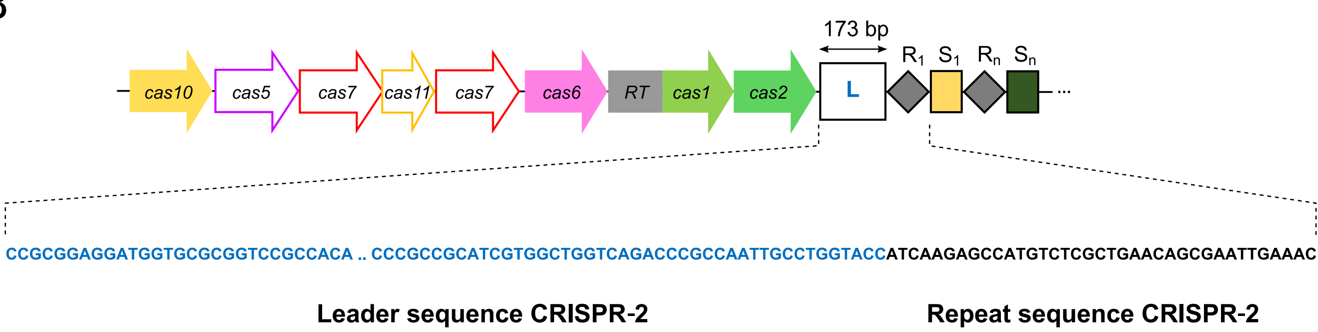


Supplementary figure S4

### Supplementary Fig S5

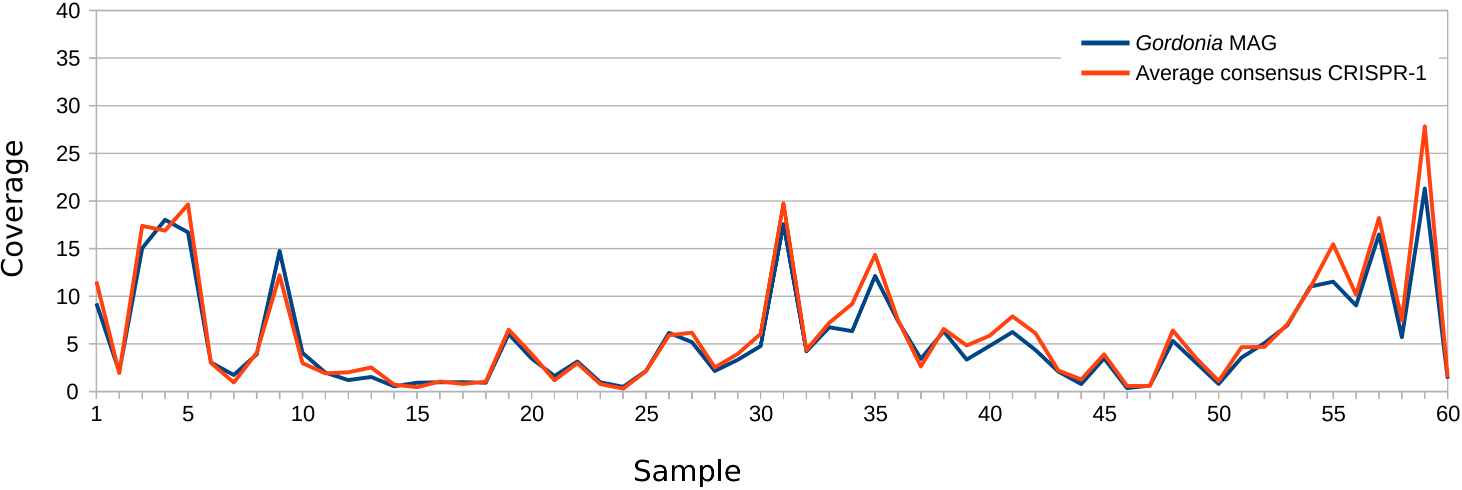


Supplementary figure S5

### Supplementary Fig S6

A

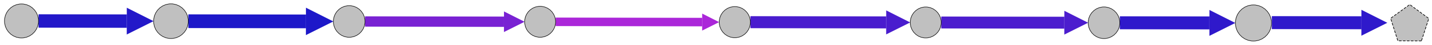

B

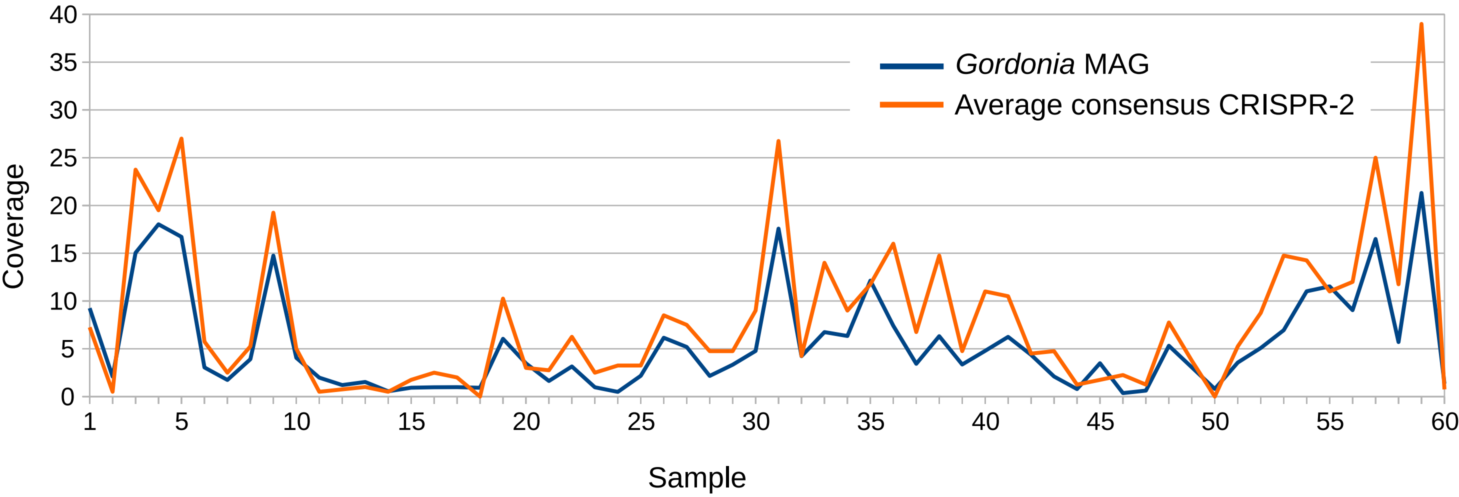


Supplementary figure S6

### Supplementary Fig S7

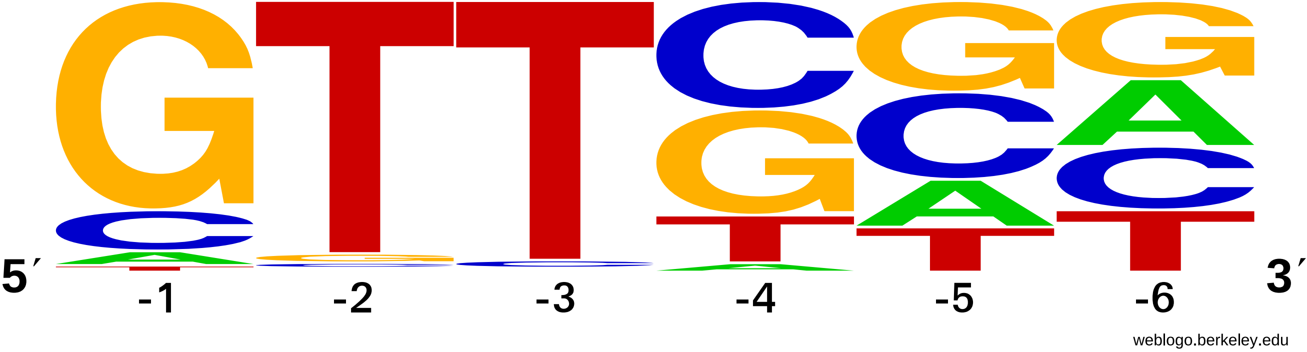


Supplementary figure S7

### Supplementary Fig S8

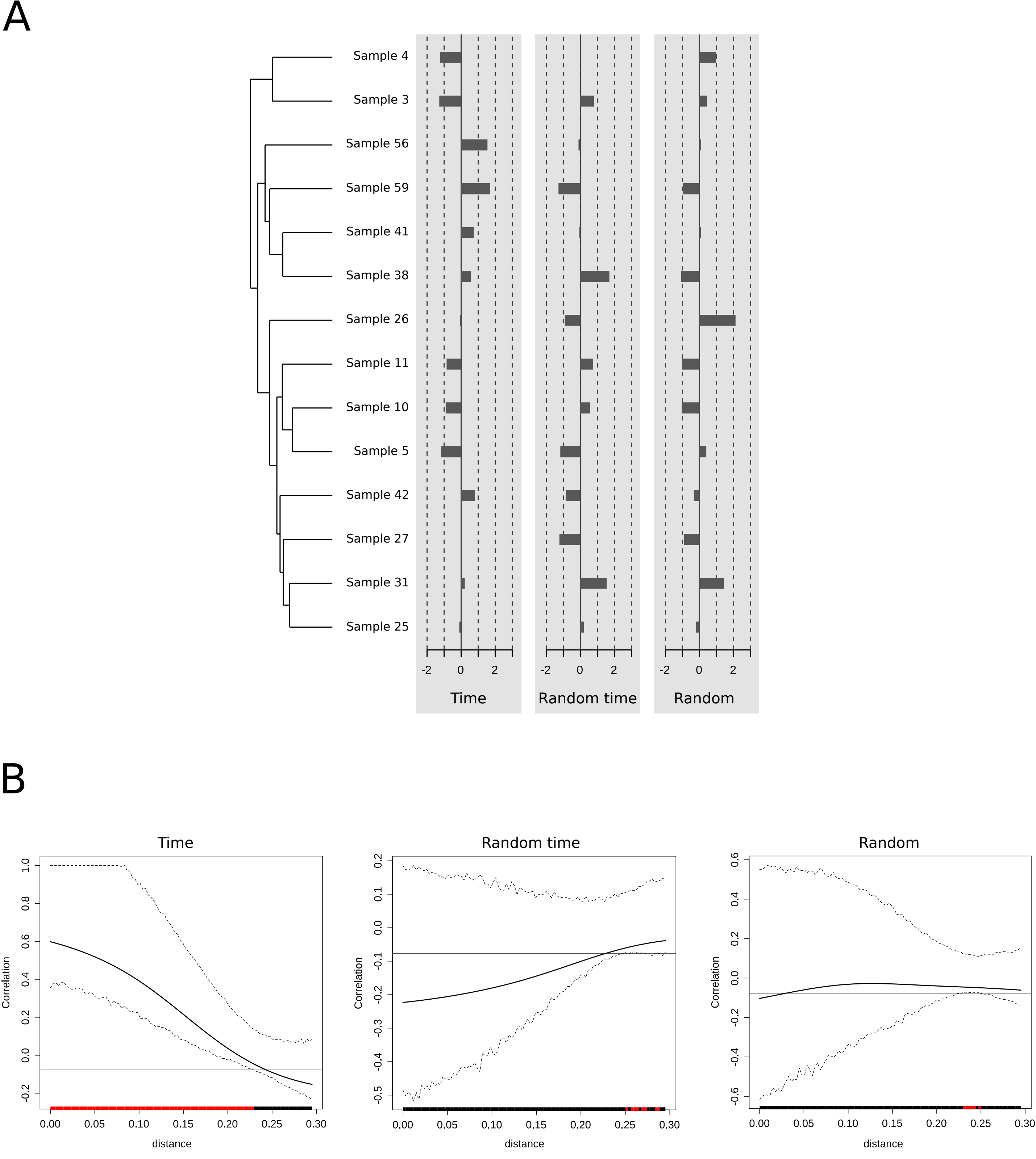


Supplementary figure S8
