## Supplementary Table S1 for "Long-run bacteria-phage coexistence dynamics under natural habitat conditions in an environmental biotechnology system"

Supplementary table 1: Pairwise ANI of *Gordonia* metagenome assembled genome compared to all available Gordonia genomes on NCBI (at March 2020)

| Name | NCBI Accesion name | Genome size (Mbp) | % GC | ANI | Alignment Fraction |
|---|---|---|---|---|---|
| Gordonia amarae NBRC 15530 | ASM24134v2 | 5.31 | 67.00 | 85.31 | 0.72 |
| Gordonia sp. NB4-1Y | ASM34729v2 | 5.63 | 65.66 | 77.73 | 0.46 |
| Gordonia rubripertincta | 53990_A01 | 4.51 | 68.50 | 77.73 | 0.44 |
| Gordonia desulfuricans | ASM1011947v1 | 5.55 | 65.58 | 77.69 | 0.47 |
| Gordonia paraffinivorans | ASM312131v1 | 4.74 | 66.09 | 77.68 | 0.45 |
| Gordonia desulfuricans NBRC 100010 | ASM148549v1 | 5.43 | 67.08 | 77.67 | 0.47 |
| Gordonia paraffinivorans NBRC 108238 | ASM34415v1 | 4.63 | 66.53 | 77.65 | 0.45 |
| Gordonia rubripertincta | ASM748926v1 | 4.86 | 65.06 | 77.53 | 0.45 |
| Gordonia rubripertincta NBRC 101908 | ASM32732v1 | 5.20 | 65.13 | 77.48 | 0.45 |
| Gordonia westfalica | IMG-taxon_2636416072_annotated_assembly | 6.41 | 63.36 | 77.47 | 0.47 |
| Gordonia rubripertincta | ASM356862v1 | 5.33 | 65.75 | 77.46 | 0.47 |
| Gordonia westfalica J30 | ASM812511v1 | 5.28 | 66.99 | 77.41 | 0.45 |
| Gordonia sp. YY1 | ASM987231v1 | 5.13 | 64.13 | 77.41 | 0.45 |
| Gordonia alkanivorans CGMCC 6845 | Goralkv1.0 | 5.03 | 64.74 | 77.37 | 0.44 |
| Gordonia bronchialis DSM 43247 | ASM2478v1 | 5.29 | 66.10 | 77.36 | 0.46 |
| Gordonia bronchialis | ASM973043v1 | 5.30 | 66.10 | 77.35 | 0.46 |
| Gordonia bronchialis | 50279_D02 | 5.33 | 65.62 | 77.35 | 0.46 |
| Gordonia alkanivorans | ASM401190v1 | 4.98 | 66.22 | 77.33 | 0.45 |
| Gordonia alkanivorans NBRC 16433 | ASM22550v1 | 5.07 | 65.55 | 77.31 | 0.45 |
| Gordonia amicalis NBRC 100051 = JCM 11271 | ASM33299v1 | 4.92 | 66.11 | 77.31 | 0.44 |
| Gordonia sp. CNJ-863 | ASM194232v1 | 5.40 | 65.17 | 77.3 | 0.44 |
| Gordonia sp. 1D | ASM232712v1 | 5.15 | 67.33 | 77.29 | 0.45 |
| Gordonia sp. AMA120 | ASM419320v1 | 7.33 | 55.75 | 77.29 | 0.46 |
| Gordonia sp. 135 | ASM972018v1 | 5.20 | 66.01 | 77.29 | 0.45 |
| Gordonia terrae C-6 | C-6 | 5.17 | 65.34 | 77.27 | 0.45 |
| Gordonia amicalis CCMA-559 | Gordonia_amicalis_CCMA-559 | 5.18 | 65.10 | 77.24 | 0.45 |
| Gordonia alkanivorans s104 | Gordonia_alkanivorans_S104 | 5.18 | 65.69 | 77.24 | 0.45 |
| Gordonia rhizosphera NBRC 16068 | ASM29819v1 | 6.43 | 66.26 | 77.23 | 0.46 |
| Gordonia namibiensis NBRC 108229 | ASM29823v1 | 4.94 | 67.44 | 77.22 | 0.45 |
| Gordonia sp. HS-NH1 | ASM141876v1 | 5.43 | 64.47 | 77.19 | 0.46 |
| Gordonia terrae | ASM284786v1 | 5.70 | 65.22 | 77.15 | 0.45 |
| Gordonia sp. UCD-TK1 | ASM169143v1 | 5.46 | 64.11 | 77.12 | 0.44 |
| Gordonia sp. ALPHA2A | ASM550260v1 | 5.19 | 67.20 | 77.12 | 0.44 |
| Gordonia sp. ALPHA1B1 | ASM550261v1 | 5.20 | 66.52 | 77.12 | 0.44 |
| Gordonia terrae | ASM550272v1 | 5.19 | 67.27 | 77.12 | 0.44 |
| Gordonia sp. JH63 | ASM985664v1 | 5.34 | 66.68 | 77.12 | 0.45 |
| Gordonia sp. IITR100 | ASM200964v1 | 5.29 | 65.13 | 77.11 | 0.44 |
| Gordonia sp. SGD-V-85 | ASM145690v1 | 5.44 | 66.41 | 77.1 | 0.45 |
| Gordonia sp. v-85 | IMG-taxon_2615840623_annotated_assembly | 5.44 | 66.41 | 77.1 | 0.45 |
| Gordonia sp. GAMMA | ASM550253v1 | 5.34 | 67.82 | 77.09 | 0.44 |
| Gordonia sp. LAM0048 | ASM165962v1 | 5.65 | 65.91 | 77.08 | 0.44 |
| Gordonia insulae | ASM385509v1 | 5.96 | 67.36 | 77.06 | 0.49 |
| Gordonia sp. HNM0687 | ASM982819v1 | 5.37 | 64.55 | 77.06 | 0.45 |
| Gordonia polyisoprenivorans HW436 | ASM38535v1 | 6.33 | 66.87 | 77.03 | 0.47 |
| Gordonia polyisoprenivorans NBRC 16320 = JCM 10675 | ASM24132v2 | 6.29 | 64.93 | 77 | 0.48 |
| Gordonia terrae NBRC 100016 | ASM24803v2 | 5.67 | 65.99 | 77 | 0.45 |
| Gordonia lacunae | ASM214901v1 | 5.76 | 66.82 | 76.99 | 0.47 |
| Gordonia terrae | ASM71697v1 | 5.67 | 65.13 | 76.98 | 0.46 |
| Gordonia terrae | ASM318382v1 | 5.71 | 67.80 | 76.98 | 0.46 |
| Gordonia terrae | 45296_E02 | 5.71 | 57.98 | 76.98 | 0.46 |
| Gordonia sp. KTR9 | ASM14388v2 | 5.89 | 65.07 | 76.96 | 0.46 |

| | | | | | |
|---|---|---|---|---|---|
| Gordonia terrae | ASM169822v1 | 5.70 | 67.81 | 76.96 | 0.46 |
| Gordonia sp. i37 | ASM204308v1 | 6.23 | 66.37 | 76.94 | 0.46 |
| Gordonia polyisoprenivorans | ASM225774v1 | 6.18 | 66.76 | 76.91 | 0.49 |
| Gordonia polyisoprenivorans VH2 | ASM24771v1 | 5.84 | 66.38 | 76.82 | 0.46 |
| Gordonia sp. SID5947 | ASM986278v1 | 5.10 | 65.00 | 76.76 | 0.44 |
| Gordonia soli NBRC 108243 | ASM33445v1 | 5.38 | 67.26 | 76.66 | 0.44 |
| Gordonia sp. RS15-1S | ASM379718v1 | 4.52 | 65.64 | 76.52 | 0.42 |
| Gordonia sp. 852002-50816_SCH5313054-a | ASM166574v1 | 5.14 | 64.73 | 76.37 | 0.43 |
| Gordonia sp. 852002-50816_SCH5313054-c | ASM166592v1 | 5.11 | 64.16 | 76.37 | 0.43 |
| Gordonia sp. 852002-50395_SCH5434458 | ASM166590v1 | 5.00 | 64.57 | 76.19 | 0.42 |
| Gordonia sihwensis | ASM96046v1 | 4.16 | 68.50 | 76.18 | 0.35 |
| Gordonia sp. 852002-10350_SCH5691597 | ASM166571v1 | 5.11 | 64.93 | 76.18 | 0.42 |
| Gordonia jacobaea | ASM118636v1 | 4.93 | 62.98 | 76.17 | 0.42 |
| Gordonia sputi NBRC 100414 | ASM24805v2 | 4.95 | 64.43 | 76.15 | 0.42 |
| Gordonia otitidis NBRC 100426 | ASM24807v2 | 5.30 | 64.95 | 76.15 | 0.41 |
| Gordonia sp. 852002-51296_SCH5728562-b | ASM166547v1 | 5.14 | 64.62 | 76.11 | 0.43 |
| Gordonia sp. NBRC 107697 | ASM993243v1 | 3.78 | 68.98 | 76.07 | 0.35 |
| Gordonia aichiensis NBRC 108223 | ASM33297v1 | 5.09 | 63.68 | 75.9 | 0.43 |
| Gordonia phthalatica | ASM130567v1 | 4.43 | 68.00 | 75.86 | 0.35 |
| Gordonia araii NBRC 100433 | ASM24126v2 | 3.91 | 66.16 | 75.85 | 0.35 |
| Gordonia sp. QH-12 | ASM157867v1 | 3.90 | 67.21 | 75.8 | 0.35 |
| Gordonia shandongensis DSM 45094 | ASM42302v1 | 3.33 | 65.79 | 75.75 | 0.32 |
| Gordonia sp. YC-JH1 | ASM284844v1 | 4.19 | 66.16 | 75.64 | 0.35 |
| Gordonia sp. NBRC 107696 | ASM993247v1 | 4.89 | 68.17 | 75.62 | 0.36 |
| Gordonia sihwensis NBRC 108236 | ASM33303v1 | 4.14 | 66.65 | 75.61 | 0.36 |
| Gordonia iterans | ASM299328v1 | 4.01 | 68.96 | 75.59 | 0.37 |
| Gordonia neofelifaecis NRRL B-59395 | ASM19243v1 | 4.26 | 67.07 | 75.56 | 0.36 |
| Gordonia hirsuta DSM 44140 = NBRC 16056 | ASM33301v1 | 3.49 | 68.35 | 75.47 | 0.34 |
| Gordonia hirsuta DSM 44140 = NBRC 16056 | ASM42068v1 | 3.49 | 68.35 | 75.47 | 0.34 |
| Gordonia kroppenstedtii DSM 45133 | ASM38048v1 | 4.23 | 64.72 | 75.39 | 0.32 |
| Gordonia hydrophobica NBRC 16057 | ASM159236v1 | 4.58 | 64.56 | 75.36 | 0.36 |
| Gordonia sp. HY186 | ASM755900v1 | 3.23 | 66.95 | 75.18 | 0.33 |
| Gordonia sp. HY189 | ASM435305v1 | 3.28 | 66.95 | 75.08 | 0.33 |
| Gordonia malaquae NBRC 108250 | ASM34413v1 | 4.47 | 64.13 | 74.98 | 0.35 |
| Gordonia malaquae | IMG-taxon_2639762576_annotated_assembly | 4.71 | 63.83 | 74.98 | 0.35 |
| Gordonia effusa NBRC 100432 | ASM24130v2 | 4.70 | 62.28 | 74.65 | 0.36 |
