## Supplementary Table S2 for "Long-run bacteria-phage coexistence dynamics under natural habitat conditions in an environmental biotechnology system"

**Supplementary Table S2**: Pearson correlation between the abundance of phage DC-56 and the abundance of the different CRISPR-1 variants over the time span of the study

| **Correlation (Pearson)** | **df[1]** | **r** | **p** |
|---|---|---|---|
| Phage DC-56 vs L1 | 58 | 0.5 | **$4.10^{-5}$** |
| Phage DC-56 vs B1 | 58 | 0.12 | 0.345 |
| Phage DC-56 vs B2 | 58 | 0.33 | **0.009** |
| Phage DC-56 vs B3 | 58 | 0.15 | 0.241 |
| Phage DC-56 vs B4 | 58 | 0.03 | 0.837 |
| Phage DC-56 vs B5 | 58 | 0.18 | 0.179 |
| Phage DC-56 vs B6 | 58 | 0.07 | 0.612 |
| Phage DC-56 vs B7 | 58 | -0.12 | 0.382 |
| Phage DC-56 vs B8 | 58 | -0.01 | 0.914 |
| Phage DC-56 vs B9 | 58 | 0.36 | **0.005** |
| Phage DC-56 vs B10 | 58 | 0.25 | 0.049 |

[1]df = Degrees of freedom
